# The pan-epigenome of the symbiotic nitrogen fixing bacterium *Sinorhizobium meliloti* unravels unexpected variability of DNA-methylation profiles in closely related strains

**DOI:** 10.1101/2023.05.31.542881

**Authors:** Iacopo Passeri, Lisa Cangioli, Marco Fondi, Alessio Mengoni, Camilla Fagorzi

## Abstract

In prokaryotes, DNA methylation has been found to be involved in several mechanisms, such as DNA repair, DNA–protein interactions, gene expression, cell cycle progression and self-DNA recognition (the Restriction-Modification systems). Studies on representatives from the same bacterial species have found that genome-wide DNA methylation patterns can be highly variable and may affect phenotypic variation and gene transfer among closely related strains. However, broader evolutionary studies on such epigenomic variation in bacteria are still scarce.

Here, we addressed this point by performing an epigenomic analysis on 21 strains of the facultative plant symbiotic nitrogen-fixing alphaproteobacterium *Sinorhizobium meliloti*. Strains of these species are characterized by a divided (multipartite) genome structure, including a chromosome, a chromid and a (more recently acquired) megaplasmid. Since these strains display extensive genomic and phenotypic variation, they are good models to test evolutionary hypotheses on the relationships among epigenomic signatures, genome structure evolution and phenotypic switches.

Results showed the presence of a wide pan-epigenome with 16 DNA methylated motifs, including both 4mC and 6mA palindromic and nonpalindromic motifs. While 9 motifs have been found methylated by all strains, the remaining had differential methylation between *S. meliloti* strains, constituting a dispensable epigenome. Differences in frequency of methylation were found among replicons, with the megaplasmid and the additional plasmids displaying several motifs with different methylation frequency with respect to the chromosome and the chromid. Moreover, differences between coding, upstream and intergenic regions, were found, suggesting that DNA methylation at specific motifs may play a role in gene regulation and consequently in phenotypic variability among strains.

Altogether, our data indicate the presence of a large epigenomic diversity in *S. meliloti*, with epigenome signatures differing between replicons, reflecting their timing of evolutionary acquisition in *S. meliloti* genomes and suggesting a role of DNA methylation in the variability of gene expression among strains.

## Introduction

Epigenetic information, namely DNA methylation and post-translational modification of histones, has been shown to be pivotal in the control of several biological phenomena in eukaryotes, such as cellular differentiation, development and pathogenesis (Jones 2012). In prokaryotic microorganisms, DNA methylation has been found involved in many molecular mechanisms such as DNA repair, DNA–protein interactions, gene expression, cell cycle progression and self-DNA recognition (the <u>R</u>estriction-<u>M</u>odification systems) (Sánchez-Romero and Casadesús 2020). Novel information gained in recent years on methylated DNA in prokaryotes, especially on Bacteria, is delving into the roles of transcriptional regulation and the formation of phenotypic variants (Oliveira and Fang 2021) (Vasu and Nagaraja 2013). Indeed, N^6^- methyladenine (6mA) is known to be involved in many events, spanning from chromosome replication to mismatch repair, conjugal transfer, antibiotic resistance (Sánchez-Romero and Casadesús 2020) bacterial differentiation (diCenzo et al. 2022) and phenotypic plasticity as phase variation (Atack et al. 2018). Besides, stress response and drug transport have been related to the presence of C^5^-methyl-cytosine (5mC))(Kahramanoglou et al. 2012; Militello et al. 2014), as N^4^-methyl-cytosine (4mC) to pathogenesis (Kumar et al. 2018). Evidence is now supporting the idea that DNA methylation in bacteria may influences phenotypes as virulence and host colonization (Blow et al. 2016; Beaulaurier et al. 2019; Oliveira 2021).

Population studies have shown large differences in R-M systems in clinical isolates of *Staphylococcus epidermidis* and in strains of the plant growth promoting bacterium (PGPB) *Bacillus velezensis* (Reva et al. 2019), where some methylation profiles were found abundant upstream to coding sequences, suggesting they can have regulatory effects on transcription. A recent evolutionary study conducted on *Helicobacter pylori* showed that phase-variable methyltransferase (MTase) genes contribute methylome variation among isolates (Estibariz et al. 2020), indicating that bacterial phenotypic variation and adaptation to variable environmental conditions may rely on epigenetic modification too. Other studies have shown that the genomic levels of 5mC relate to antibiotic resistance (Militello et al. 2014; Yuan et al. 2021). More recently, a pan-epigenome analysis of *Mycoplasma agalactiae* revealed the existence of strain-specific motifs, which allowed the identification of orphan MTases (Dordet-Frisoni et al. 2022), affecting the rate of horizontal gene transfer events among *M. agalactiae* strains. Consequently, it is becoming highly relevant for molecular evolutionary studies in bacteria to investigate the extent of the pan-epigenome and evaluate its role in genome evolution and adaptation.

Facultative symbiotic bacteria are relevant models for the study of environmental adaptation, since they colonize multiple environments coping with dramatically contrasting selective pressure. *Sinorhizobium meliloti* is a facultative plant symbiotic bacterium, belonging to the *Rhizobiaceae* family, able to thrive on soil, colonizing the surrounding of plant roots and entering in symbiosis with host plants (mainly from the Leguminous genera *Medicago, Melilotus* and *Trigonella*), where it differentiates into bacteroid, establishing an intracellular symbiosis in specialized root organs called nodules and fixing atmospheric dinitrogen into ammonia (Poole et al. 2018; Cangioli et al. 2021; Ledermann et al. 2021). *S. meliloti* is one of the most studied species for what is concerning symbiotic nitrogen fixation (diCenzo et al. 2019). The genomes of several *S. meliloti* strains have been sequenced and many population-genomic and evolutionary studies have been performed on this species in relation to adaptation and selective pressure for symbiosis (see for instance (Batstone et al. 2020; Batstone et al. 2022; Epstein et al. 2022) and multipartite genome evolution (Galardini, et al. 2013; diCenzo et al. 2016). In fact, the strains of this species often harbour a multipartite (or divided) genome, as for many bacterial species interacting with hosts (diCenzo and Finan 2017; diCenzo, et al. 2019), composed by three main replicons: a chromosome, a chromid and a megaplasmid. The latter contains the essential machinery for symbiosis, while the other two are mainly relevant for growth in soil and in the plant rhizosphere (diCenzo et al. 2016 and 2014). As in the other alphaproteobacterial species (Zweiger et al. 1994), *S. meliloti* cell cycle is controlled by the action of the master regulator CtrA, which in turn expresses its activity in relation to DNA methylation status on a pentanucleotide sequence (GANTC), methylated by the orphan DNA MTase CcrM (Fioravanti et al. 2013; Pini et al. 2015). Recently, we investigated the profile of DNA methylation in two strains of *S. meliloti* along the intracellular differentiation phases in the plant root nodule (diCenzo et al. 2022). In this study, we reported evidence that the CcrM activity on GANTC sequence is dysregulated during symbiosis, further suggesting this dysregulation is a driving factor for intracellular differentiation of *S. meliloti* cells into bacteroids. In this work we noticed the presence of methylated motifs other than GANTC sequence motifs. These additional motifs were found with lower occurrence than GANTC, but interestingly many of them were present in only one of the investigated *S. meliloti* strains, suggesting the presence of a pan-epigenome, composed by a shared set of methylated motifs plus a few unshared or strain-specific motifs. In agreement with the presence of a pan-epigenome, in the same work (diCenzo et al. 2022) we analyzed 24 putative DNA MTases identified in a set of 20 *S. meliloti* genomes and we found that CcrM was the only one in common to all the strains, while many others were in single strains only. These DNA MTases present in the dispensable genome fraction can be the genetic determinants of the unshared methylated motifs (viz. dispensable epigenome). Such motifs could be possibly related to R-M systems and determine the formation of barriers to gene flow. Indeed, in a previous work we have shown for the *S. meliloti* 1021 strain, that the *hsd*R gene, coding for a putative type-I restriction enzyme, plays a role in the frequency of gene transfer from different donor strains (Ferri et al. 2010), suggesting that the observed differences in DNA methylation profiles and DNA MTases in *S. meliloti* may contribute barriers to gene flow and then to the establishment of subpopulations. However, since evidence for a role of DNA methylation in phenotypic variation in bacteria are increasing, we cannot exclude that both the core and the dispensable epigenome could impact on regulation of gene expression. *S. meliloti* strains, beside displaying relevant differences in genome content because of the typical open pangenome structure (Galardini et al. 2013), also present large phenotypic and transcriptomic variation, both in free-living growth and in symbiotic-related conditions (Biondi et al. 2009; Bellabarba et al. 2021; Fagorzi et al. 2021). We may likely hypothesize that this large variability could also be mirrored by genome-wide DNA methylation variation.

In this work, we examined genome-wide methylation pattern variations in *S. meliloti* strains to: i) assess the presence of a pan-epigenome, ii) evaluate its relationship with the pangenome structure and a possible effect on gene transfer (i.e. HGT) and multireplicon structure, and iii) inspect the occurrence of an uneven distribution of methylated motifs with respect to coding sequences, thus indicating the possibility to relate epigenomic differences to gene regulation. Results obtained showed the presence of a wide pan-epigenome, including both 4mC and 6mA palindromic and nonpalindromic motifs. These motifs have differential occurrence in *S. meliloti* replicons and between coding and noncoding sequences and partially mirroring gene exchange rates. Taken together, our results suggest that the epigenome may impact *S. meliloti* microevolution and adaptation.

## Results

### S. meliloti has a large pan-epigenome

SMRT sequencing of 21 *S. meliloti* strains yielded genomes with an average coverage of 155X, average genome size of 7.1 Mbp and from 3 to 9 contigs. Contig mapping to *S. meliloti* 1021 genome allowed to identify, for 16 strains, a certain number of contigs not mapping to the sequences of *S. meliloti* 1021 chromosome, pSymB chromid and pSymA megaplasmid. These contigs were labelled as “other” and range from few Kbp (e.g., 3’455 bp in AK75) to hundreds of Kbp (e.g., 328’353 bp for CE480L). We may hypothesize that some of these contigs could correspond to additional plasmids, as already shown for AK83 and SM11 strains (Schneiker-Bekel et al. 2011; Galardini et al. 2011) and as suggested by the number of genes coding for the replication initiation protein RepC identified in the genome sequences (Supplementary Information file, Table S1). Considering the entire panel of strains, a total of 27 methylated raw motifs were identified, further collapsed into 16 different motifs (Supplementary Information file, Table S2), since they shared a common core sequence. Seven motifs contained 4mC, while nine contained 6mA. Ten motifs were palindromic or nearly palindromic. The proportion of motifs methylated ranged from an average of 0.02 to nearly fully methylated (0.99) for GANTC (Supplementary dataset S3). The nearly full methylation for GANTC (0.98-0.99) is in line with the expectation from cells collected in the late exponential growth phase, as previously observed (diCenzo et al. 2022).

Among the identified methylated motifs, two groups can be defined: a “core” set of motifs (shared by all strains analyzed) and a “dispensable” one (occurring in some strains only). The core epigenome was composed of 9 motifs, which are methylated in all strains, while the other 7 motifs formed the dispensable epigenome, being methylated from a minimum of 2 to a maximum of 19 strains. No strain-exclusive motif was found (Figure 1, Supplementary dataset S3). The core GANTC motif is that recognized by the highly conserved cell cycle regulated CcrM methyltransferase and it was expected to be part of the core epigenome (being *ccrM* part of the core genome), as previously reported (diCenzo et al. 2022). We here hypothesize that the other core motifs are due to a common set of DNA-MTases shared between the strains, while the dispensable epigenome suggests the presence of DNA MTases encoded by genes belonging to the accessory genomes. Indeed, genome annotation identified from 4 to 9 DNA MTases in the genomes (Supplementary information file, Table S3, and Supplementary dataset S2), which agrees with the variability in methylated DNA motifs found among strains. When inspecting if strains with differences in the methylation profiles were showing differences in the transfer of genetic material too (possibly due to methylation profiles linked to R-M systems), we found significant differences (up to 2 orders of magnitude) in pairwise gene transfer efficiency in five randomly selected strains (AO641M, BL225C, CE480L, CL374FF and CO438LL) (Supplementary information file, Figure S1 and Tables S4-S6). This result may suggest that differential presence of methylated profiles could have some impact on gene transfer among strains of *S. meliloti*.

**Figure 1.**
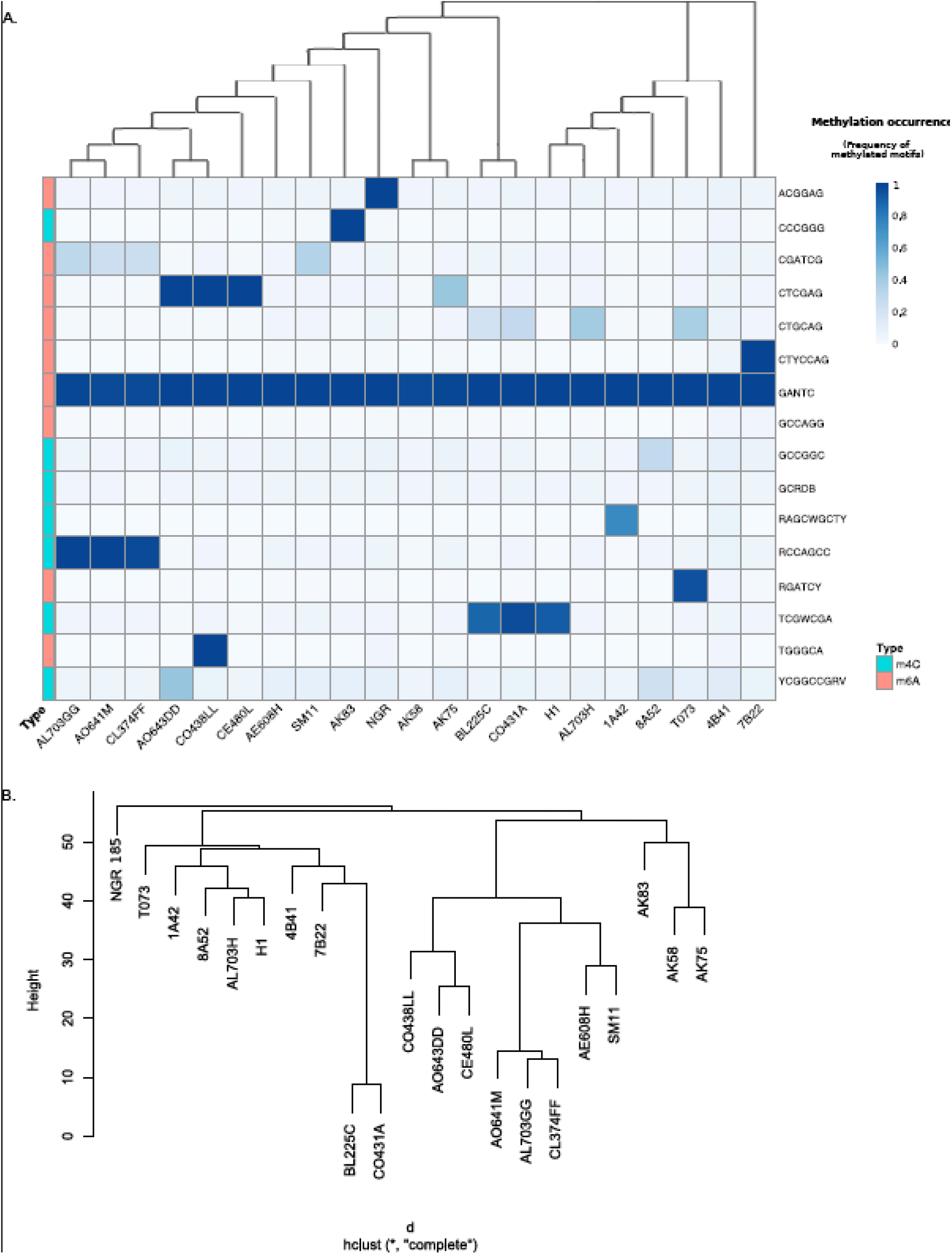
*S. meliloti* pan-epigenome partially reflects strains genomic relatedness. A) Matrix of occurrence of methylated profiles and their level of methylation. Clustering of strains is based on unrooted core gene phylogeny from a concatemer of 1’484’250 nucleotide residues. Clustering of motifs is based on Bray-Curtis distance. B) Clustering of the dispensable genome based on 11’178 genes.

Finally, the phylogenetic relationships among these strains have a relatively limited overlap with similarities in their epigenomes. Among strains whose shared epigenome pattern mirrors their phylogenetic relationships we found strains AO641M, AL703GG and CL347FF. These are evolutionarily strongly related and share a high density of methylation on RCCAGCC (Figure 1). In most of the other cases (e.g., SM11), nor core or dispensable genome phylogenies clearly reflect epigenome profiles, overall indicating that both the presence and the frequency of methylation in *S. meliloti* only partially reflects genomic relatedness.

### Intergenic and upstream sequences display low methylation values

To check whether the detected methylation profiles could be involved in gene expression control, we inspected the occurrence of methylated motifs in coding, non-coding, intergenic and upstream to genes regions (CDS, nCDS, tIG and US sequences, as defined in Figure S2) in each of the 21 genomes analyzed (Supplementary Dataset S3). In general, tIG sequences showed a statistically significant lower frequency of methylated motifs with respect to CDS, nCDS and US for most of the motifs (p<0.05 one-way ANOVA, Dunn’s post hoc test, Supplementary Dataset S3). These results indicate that methylation frequency is not evenly distributed across functional genomic features (i.e., CDS, tIG, US). Methylation of specific DNA sequence motifs is known to be related to DNA transcriptional regulation (Sanchez-Romero and Casadesus 2020). For instance, in *Caulobacter crescentus* the binding properties and activation of gene expression by the transcription factor GcrA is dependent on the methylation of the coding DNA strains (Mohapatra et al. 2020). The lower level of methylation in tIG may suggest possible role in transcriptional regulation of some of the found motifs. The biplot of a Principal Component analysis of the methylation frequencies across motifs (Figure 2b) reinforced this hypothesis, clearly showing a genomic feature-dependent clustering and a strong separation of regions likely containing putative regulatory elements (tIG and US) from coding sequences (CDS and nCDS) of the variance vectors. To evaluate the strain-by-strain variation across functional features, the coefficient of variation (CV) in frequency of methylation was computed (Figure 2c). The smallest values of CV, corresponding to the lowest inter-strain variability were found for the highly abundant methylated motifs GANTC and GCRDB, and for GCCAGG (ranging between 10^−5^ and 10^−3^). The other motifs exhibited a CV of methylated fractions 100 times higher than that of these highly methylated motifs (ranging around 10^−1^), which may further suggest an involvement of methylation of such motifs in methylation-mediated variability of phenotype/gene expression among strains. However, additional investigation of methylation pattern under different growth conditions coupled with transcriptomic analyses are needed to sustain the hypothesis of an epigenomic cause of phenotypic plasticity and transcriptional variation in *S. meliloti* strains.

**Figure 2.**
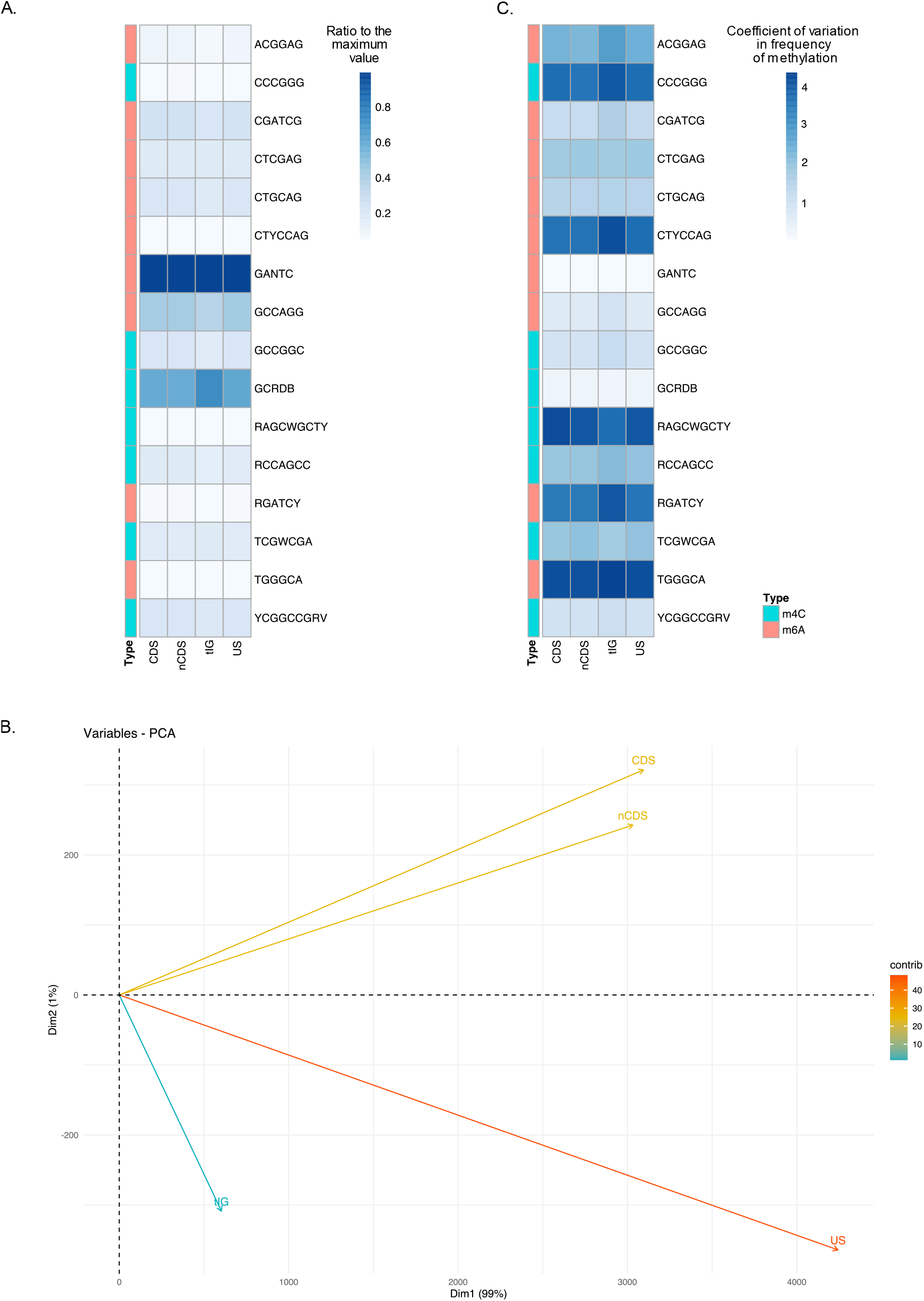
Functional genomic features show different methylation status. A) Heatmap of the proportion of ratio of methylation (ratio to the maximal value) functional features; B) Principal Component Analysis of values of frequency of methylation for tIG, US, CDS and nCDS. A biplot reporting the motifs mostly contributing to differences among features is reported; C) Heatmap of the coefficient of variation in frequency of methylation.

### Methylation profiles are mainly similar across replicons

Since *S. meliloti* harbors a multipartite genome structure, which is connected to a functional specialization of the different replicons (chromosome, a chromid and a megaplasmid) and to different evolutionary dynamics (Galardini et al. 2013; diCenzo et al. 2014; diCenzo et al. 2016), we tested the following hypothesis: can DNA methylation profiles reflect such intragenomic differences among replicons? The differential density of methylated motifs along the multipartite *S. melilot*i genome was then analyzed, to check for the presence of a possible uneven occurrence of methylated motifs across the *S. meliloti* replicons (chromosome, pSymB chromid, pSymA megaplasmid and “other”) (Figure 3; Supplementary Dataset 3). Out of the 16 motifs identified, 10 showed differences in the frequency of methylation between “other” and the rest of the replicons, 4 showed differences between the chromosome and pSymA, 4 between pSymA and pSymB, and 1 only between the chromosome and pSymB. Interestingly, the methylation frequencies of “other” replicons displayed differences even for GANTC motifs (and particularly low in AK83, 0.81 vs. 0.99 in the rest of the genome), suggesting an uncoupling of the control of DNA duplication of such elements (likely corresponding to plasmids, as indicated by the number of *repC* genes found, Supplementary information file). This possible uncoupling of DNA replication from cell cycle control in the divided *S. meliloti* genome was previously suggested for the megaplasmid pSymA in the terminally differentiated cells (bacteroids) inside root nodules (diCenzo et al. 2022). Moreover, since the “other” replicons are part of the dispensable genome fraction, we cannot exclude that the observed differential frequency of some methylation motifs may reflect their recent HGT. Indeed, it has been shown in rhizobia that genes deriving from recent horizontal gene transfer (HGT) events can be subjected to transcriptional silencing (Shi et al. 2022).

**Figure 3.**
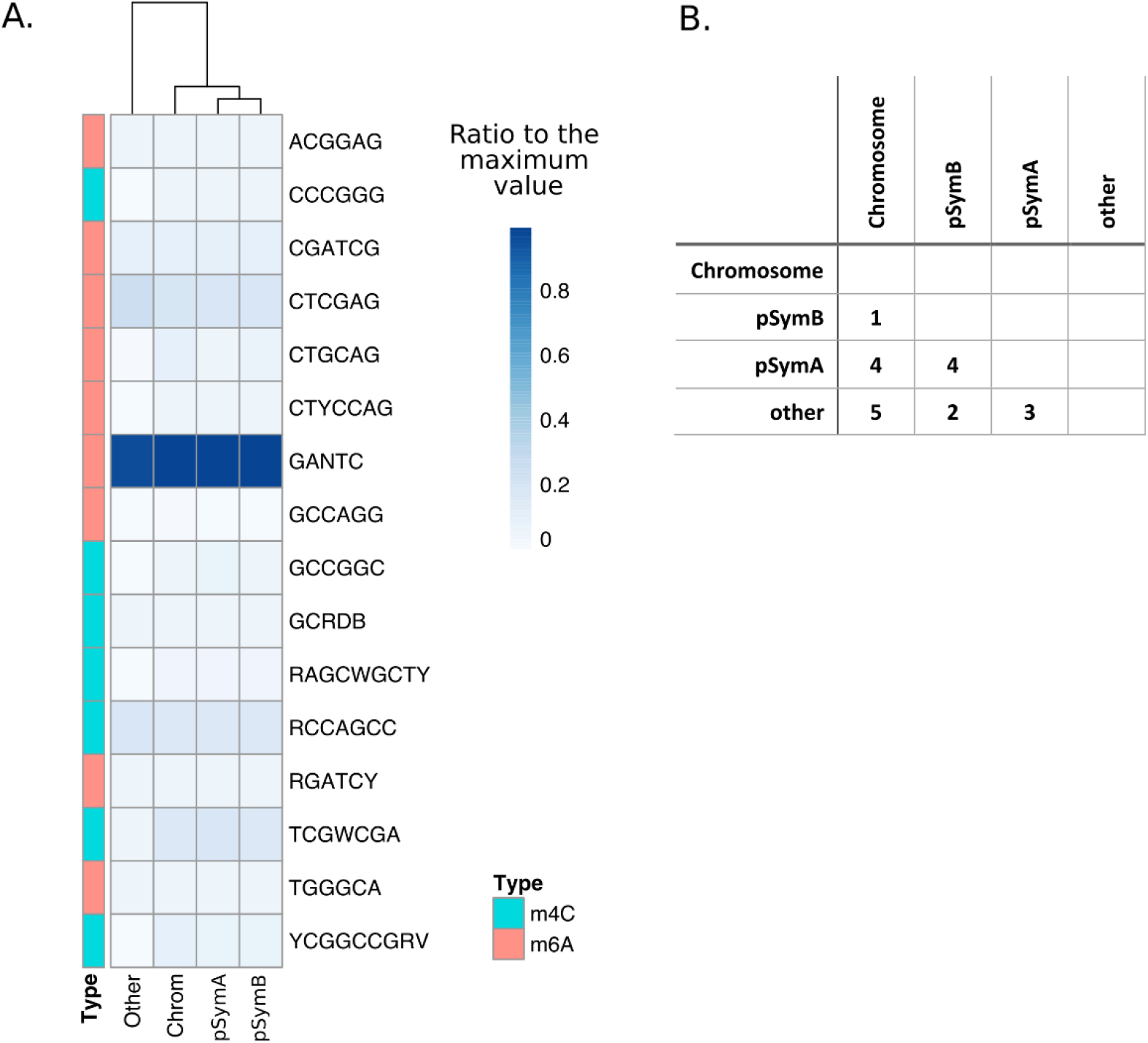
Relative frequency of methylated motifs across the multipartite replicons of *S. meliloti*. A) Heatmap reporting the ratios to maximum values for each motif. B) Number of methylated motifs with statistically significant differences among replicons.

## Discussion

Genome-wide DNA methylation patterns are key elements in the genotype-to-phenotypes translations, having profound consequences on adaptation and phenotypic plasticity (Herman and Sultan 2016). In prokaryotes, DNA methylation can have several roles, from DNA repair mechanisms, cell cycle control, self-DNA recognition and transcriptional regulation (Oliveira and Fang 2021). Mirroring these multiple roles of DNA methylation, the phylogeny of DNA methyltransferases is also complex, including both vertical and horizontal transmission (Harris and Goldman 2020). This, in turn, may give rise to both DNA methylation patterns shared among strains and to strain-specific DNA methylation profiles, resulting from genes harbored in the core and in the dispensable genome fraction, respectively. In agreement with such hypotheses, evidence from our data on *S. meliloti* indicates that a shared by all (core) and an unshared (dispensable) set of DNA methylation profiles exist. Such epigenomic landscape, with a core and dispensable epigenome, is in line with the pangenomic structure of this species (Galardini et al. 2013). However, the phylogenetic trees on core and dispensable genomes of the *S. meliloti* strains only partially mirror epigenome similarities, suggesting that epigenome profiling (presence and frequency of methylated DNA motifs) cannot be simply explained by evolutionary relatedness. Indeed, considering the possibility that a substantial fraction of the methylated motifs may derive from HGT events (being the motifs occurring in a part only of the strains) we should consider the differential influence acquired DNA methylation profiles may have on the following acquisitions of DNA. When DNA methylation profiles are linked to R-M systems, their acquisition strongly reduces the acquisition of novel DNA (and of potentially novel DNA methyltransferase and consequently new DNA methylation profiles) (Jeltsch 2003). This may bring to a reduction of gene flow among strains with different methylation profiles, favoring the maintenance of cohesive population structures, which in turn can give rise to a new lineage in the population (Oliveira et al. 2016). On the contrary, acquired DNA methylation profiles unlinked to the restriction systems have potentially no effect over novel acquisitions. Consequently, we can figure out that the pattern of the dispensable (viz. unshared) epigenome is the result of the timing of acquisition of “neutral” and “negative” (to the HGT) profiles, corresponding to orphan DNA MTases and R-M systems, respectively. This model will lead to a mirroring of the dispensable genome phylogeny only for those branches where mostly “neutral” profiles have been acquired. The experimental evidence of differences among strains for gene transfer frequencies in our study may suggest the presence of some “negative”-to- HGT methylation profiles. However, from our data we cannot rule out which profile can be more related to differences in gene transfer among the selected strains.

Still concerning the relationships between HGT and DNA methylation, we inspected the multi-replicon structure of *S. meliloti* genome with respect to the pattern of methylation of the chromosome, the chromid pSymB and the symbiotic megaplasmid pSymA, the three resident replicons of *S. meliloti* (Galibert et al. 2001). Evolutionary analyses on *S. meliloti* genomes clearly indicated that pSymA megaplasmid is an alien element which was recently acquired by the resident genome, while pSymB is a chromid, which stabilizes in the ancestral *S. meliloti* genome (diCenzo et al. 2014; Galardini et al. 2013, Galibeert et al. 2001). In line with this timing of replicon acquisition, statistically significant differences between the chromosome and the other replicons were limited for pSymB (1 motif only), but higher for pSymA (4 motifs) and “other” (7 motifs). As expected, differences between pSymA and pSymB were relevant and in line with those between pSymA and the chromosome (4 motifs). This evidence possibly indicates that the DNA methylation status is another parameter which can allow to define genomic integration in multipartite genomes, as differences between chromosome, chromid and plasmids. In line with such results, there are recent indication that HGT elements (i.e., plasmids) show avoidance of target of R-M systems, which may be reflected in different methylation abundance for specific target motifs (Shi et al. 2022)). Moreover, DNA regions originating from HGT, in general tend to have low expression levels and slowly integrate into the transcriptional network (Lercher and Pál 2008). This low level of expression can be related to the presence of xenogeneic silencers, as MucR in *Sinorhizobium fredii* (Jiao et al. 2022), which preferentially targets regions with high AT%. We may speculate that the different methylation frequency for some motifs of “other” replicons may partially reflect a stronger control of gene expression operated during the growth conditions applied.

A hypothesis on the role of the DNA methylation on gene expression regulation was drawn from the results obtained when measuring the frequency of methylation in upstream, intergenic and coding sequences where intergenic regions showed for some motifs lower values of methylation frequencies compared to coding sequences, as well as a high strain-by-strain variability. Such results in fact can be interpreted in terms of involvement of DNA methylation in gene expression control operated at the promoter/intergenic level by differential binding/accessibility of transcription factors. Large strain-by-strain variation in transcriptomes of *S. meliloti* strains has been previously shown and related to genotypic interaction with host plant (Fagorzi et al. 2021). We cannot a priori exclude that strain-by-strain epigenomic differences can have an influence over the observed transcriptomic variations.

In conclusion, here we showed for the first time that, in bacteria, a large pan-epigenome can exist. Moreover, using *S. meliloti* (a nitrogen fixing, multipartite genome model organism), while highlighting that replicons share a relatively similar methylation profiles, we recognized that upstream and intergenic regions can be less methylated than coding sequences and that this pattern of differential methylation is highly variable among strains. These data allow to sustain the hypothesis that DNA methylation may play a role in gene regulation and in phenotypic variation at the strain level. Investigation on DNA MTases mutants and additional analyses of genome-wide methylation in different culture conditions are needed for thoroughly testing this hypothesis.

## Materials and Methods

### S. meliloti strains, growth conditions and DNA extraction

*S. meliloti* strains used in this work are listed in Table S7 (Supplementary Information File). All strains were routinely grown on TY medium. Overnight cultures of all strains grown in 10 ml TY medium overnight at 30 ° C with shaking (130 rpm). To obtain late exponential/stationary phase samples, cultures were harvested after 24 hours of growth at OD_600nm_ values of ∼ 1.4. In all cases, cultures were streaked on TY plates to check for contamination and then centrifuged (8,200 g, 10 minutes, 4°C); the full 10 mL was centrifuged and transferred to a 2 mL tube. DNA was isolated using Qiagen PowerLyser PowerSoil Pro KIT according to manufacturer instructions.

### SMRT genome sequencing of S. meliloti strains

Single-Molecule Real Time (SMRT) sequencing was performed in house at the University of Florence using Pacific Biosciences Sequel instrument (Eid et al. 2009). Genomic DNA was sheared to 20 Kbp using g-TUBEs (Covaris Inc., Woburn, MA, USA). Sheared DNA was treated with exonuclease to remove single stranded ends and DNA damage repair mix followed by end repair and ligation of barcoded blunt adapters using SMRTbell Template Prep Kit 2.0 (PacBio, Menlo Park, CA, USA). Libraries were purified with AMPure PB beads (Beckman Coulter, Brea, CA, USA) and eight libraries with different barcodes were pooled at equimolar ratios and purified with AMPure PB beads. SMRTbell template libraries were prepared using a Sequel Binding Kit 3.0 (PacBio, Menlo Park, CA, USA), and sequenced on a Sequel instrument using a v3 or v4 sequencing primer, 1M v3 SMRT cells, and Version 3.0 sequencing chemistry.

Genome assembly was performed using the PacBio SMRT Link software (PacBio, Menlo Park, CA, USA). Briefly, raw reads were filtered using SFilter to remove short reads and reads derived from sequencing adapters. Filtered reads were assembled into contig though the Microbial Assembly SMRT Link tool. Each contig has been aligned with the genome of *S. meliloti* 1021 (GenBank assembly accession GCA_000006965.1) using Mauve software (Darling et al. 2004) and categorized into Chromosome, pSymA-like replicon, pSymB-like replicon and “other” (when no mapping to *S. meliloti* 1021 was found) (Supplementary dataset S1). Genomes were reannotated with Prokka (V 1.12-beta). Genome sequences are deposited under NCBI BioProject Accession PRJNA681719.

### Identification of methylated bases and mapping to genomic features

The assembled genomes were used as reference to map modified sites identified through kinetic analysis of the aligned DNA sequence data with the SMRT Link software ver.8.0.0.80529 (Pacific Biosciences, Menlo Park, CA, USA) (Flusberg et al. 2010), using default options; the number of mapped bases per sample is provided in Supplementary dataset S1. Modified sites were then grouped into motifs using MotifFinder. Downstream analyses were performed using MeStudio (Riccardi et al. 2022). MeStudio input files are the genome sequences in FASTA format, the genome annotation in GFF3 format and the PacBio-derived output file with methylation annotations in GFF3 format. MeStudio allows to determine the motifs distribution along: i) protein-coding gene with accordant (sense) strand (CDS), ii) regions that fall between annotated genes (true intergenic, tIG), iv) regions upstream to the reading frame of a gene, with accordant strand (US) (Supplementary information file, Figure S2). The differences in the frequencies of methylated motifs among categories were evaluated by post hoc Dunn’s test and Pearson correlation.

### Phylogenetic reconstruction

To construct an unrooted core gene phylogeny, the pangenome of the 21 *S. meliloti* strains was calculated using Roary (version 3.11.2) (Page et al. 2015) with a percent identity threshold of 95% (Supplementary dataset S2). The nucleotide sequences of the 4,633 core genes (identified as those found in at least 99% of the genomes; Supplementary dataset S2) were aligned with PRANK (Löytynoja 2014) and the alignments were trimmed and concatenated. The concatenated alignment was used to construct a maximum likelihood phylogeny (the bootstrap best tree following 100 bootstrap replicates) using RAxML 8.4.2 (Stamatakis 2006) with the GTRCAT model. Phylogeny was visualized with the online iTOL webserver (Letunic and Bork 2021).The dispensable gene set of the 21 strains was used to create a presence/absence matrix. Hierarchical clustering was performed in R (R Core Team 2014) with *hclust* function based on Euclidean distance, with “complete” agglomeration method.

## Acknowledgement

This work was partially supported by the grant MICRO4Legumes, D.M.n.89267 (Italian Ministry of Agriculture) to A.M. I.P. is supported by a PhD fellowship from PNRR, D.M. n. 351/2022 (Italian Ministry of University and Research). L.C. is supported by a PhD fellowship from MICRO4Legumes. C.F. is supported by a post-doctoral fellowship from the LEGU-MED project (PRIMA Foundation) Call 2019. We are grateful to George C. diCenzo for useful insights and suggestions during data analysis and manuscript writing.

## References

Atack JM, Yang Y, Seib KL, Zhou Y, Jennings MP. 2018. A survey of Type III restriction-modification systems reveals numerous, novel epigenetic regulators controlling phase-variable regulons; phasevarions. Nucleic Acids Res.

Batstone RT, Burghardt LT, Heath KD. 2022. Phenotypic and genomic signatures of interspecies cooperation and conflict in naturally occurring isolates of a model plant symbiont. Proceedings of the Royal Society B: Biological Sciences 289.

Batstone RT, O’Brien AM, Harrison TL, Frederickson ME. 2020. Experimental evolution makes microbes more cooperative with their local host genotype. Science (1979).

Beaulaurier J, Schadt EE, Fang G. 2019. Deciphering bacterial epigenomes using modern sequencing technologies. Nat Rev Genet.

Bellabarba A, Bacci G, Decorosi F, Aun E, Azzarello E, Remm M, Giovannetti L, Viti C, Mengoni A, Pini F. 2021. Competitiveness for Nodule Colonization in Sinorhizobium meliloti: Combined In Vitro -Tagged Strain Competition and Genome-Wide Association Analysis. mSystems.

Bertalan M, Albano R, de Pádua V, Rouws L, Rojas C, Hemerly A, Teixeira K, Schwab S, Araujo J, Oliveira A, et al. 2009. Complete genome sequence of the sugarcane nitrogen-fixing endophyte Gluconacetobacter diazotrophicus Pal5. BMC Genomics.

Biondi EG, Tatti E, Comparini D, Giuntini E, Mocali S, Giovannetti L, Bazzicalupo M, Mengoni A, Viti C. 2009. Metabolic capacity of Sinorhizobium (Ensifer) meliloti strains as determined by phenotype microarray analysis. Appl Environ Microbiol.

Blow MJ, Clark TA, Daum CG, Deutschbauer AM, Fomenkov A, Fries R, Froula J, Kang DD, Malmstrom RR, Morgan RD, et al. 2016. The Epigenomic Landscape of Prokaryotes. PLoS Genet.

Cangioli L, Checcucci A, Mengoni A, Fagorzi C. 2021. Legume tasters: Symbiotic rhizobia host preference and smart inoculant formulations. Biological Communications.

Darling ACE, Mau B, Blattner FR, Perna NT. 2004. Mauve: Multiple alignment of conserved genomic sequence with rearrangements. Genome Res.

diCenzo GC, Cangioli L, Nicoud Q, Cheng JHT, Blow MJ, Shapiro N, Woyke T, Biondi EG, Alunni B, Mengoni A, et al. 2022. DNA Methylation in Ensifer Species during Free-Living Growth and during Nitrogen-Fixing Symbiosis with Medicago spp.. mSystems.

DiCenzo G.C., Checcucci A, Bazzicalupo M, Mengoni A, Viti C, Dziewit L, Finan TM, Galardini M, Fondi M. 2016. Metabolic modelling reveals the specialization of secondary replicons for niche adaptation in Sinorhizobium meliloti. Nat Commun 7.

DiCenzo George C., Checcucci A, Bazzicalupo M, Mengoni A, Viti C, Dziewit L, Finan TM, Galardini M, Fondi M. 2016. Metabolic modelling reveals the specialization of secondary replicons for niche adaptation in Sinorhizobium meliloti. Nat Commun.

diCenzo GC, Finan TM. 2017. The Divided Bacterial Genome: Structure, Function, and Evolution. Microbiology and Molecular Biology Reviews.

diCenzo GC, MacLean AM, Milunovic B, Golding GB, Finan TM. 2014. Examination of Prokaryotic Multipartite Genome Evolution through Experimental Genome Reduction. PLoS Genet.

Dicenzo GC, Mengoni A, Perrin E. 2019. Chromids Aid Genome Expansion and Functional Diversification in the Family Burkholderiaceae. Mol Biol Evol.

Dicenzo GC, Zamani M, Checcucci A, Fondi M, Griffitts JS, Finan TM, Mengoni A. 2019. Multidisciplinary approaches for studying rhizobium–legume symbioses. Can J Microbiol.

Dordet-Frisoni E, Vandecasteele C, Contarin R, Sagné E, Baranowski E, Klopp C, Nouvel LX, Citti C. 2022. Impacts of Mycoplasma agalactiae restriction-modification systems on pan-epigenome dynamics and genome plasticity. Microb Genom 8:1–23.

Eid J, Fehr A, Gray J, Luong K, Lyle J, Otto G, Peluso P, Rank D, Baybayan P, Bettman B, et al. 2009. Real-time DNA sequencing from single polymerase molecules. Science (1979).

Epstein B, Burghardt LT, Heath KD, Grillo MA, Kostanecki A, Hämälä T, Young ND, Tiffin P. 2022. Combining GWAS and population genomic analyses to characterize coevolution in a legume-rhizobia symbiosis. Mol Ecol:1–14.

Estibariz I, Ailloud F, Woltemate S, Bunk B, Spröer C, Overmann J, Aebischer T, Meyer TF, Josenhans C, Suerbaum S. 2020. In vivo genome and methylome adaptation of CAG-negative helicobacter pylori during experimental human infection. mBio.

Fagorzi C, Bacci G, Huang R, Cangioli L, Checcucci A, Fini M, Perrin E, Natali C, diCenzo GC, Mengoni A. 2021. Nonadditive transcriptomic signatures of genotype-by-genotype interactions during the initiation of plant-rhizobium symbiosis. mSystems 6.

Ferri L, Gori A, Biondi EG, Mengoni A, Bazzicalupo M. 2010. Plasmid electroporation of Sinorhizobium strains: The role of the restriction gene hsdR in type strain Rm1021. Plasmid.

Fioravanti A, Fumeaux C, Mohapatra SS, Bompard C, Brilli M, Frandi A, Castric V, Villeret V, Viollier PH, Biondi EG. 2013. DNA Binding of the Cell Cycle Transcriptional Regulator GcrA Depends on N6-Adenosine Methylation in Caulobacter crescentus and Other Alphaproteobacteria. PLoS Genet.

Flusberg BA, Webster DR, Lee JH, Travers KJ, Olivares EC, Clark TA, Korlach J, Turner SW. 2010. Direct detection of DNA methylation during single-molecule, real-time sequencing. Nat Methods.

Galardini M, Mengoni A, Brilli M, Pini F, Fioravanti A, Lucas S, Lapidus A, Cheng JF, Goodwin L, Pitluck S, et al. 2011. Exploring the symbiotic pangenome of the nitrogen-fixing bacterium Sinorhizobium meliloti. BMC Genomics.

Galardini M, Pini F, Bazzicalupo M, Biondi Emanuele G., Mengoni A. 2013. Replicon-dependent bacterial genome evolution: The case of Sinorhizobium meliloti. Genome Biol Evol.

Galardini M, Pini F, Bazzicalupo M, Biondi Emanuele G, Mengoni A. 2013. Replicon-Dependent Bacterial Genome Evolution : The Case of. Mol Biol 5:542–558.

Galibert F, Finan TM, Long SR, Pühler A, Abola P, Ampe F, Barloy-Hubler F, Barnet MJ, Becker A, Boistard P, et al. 2001. The composite genome of the legume symbiont Sinorhizobium meliloti. Science (1979).

Harris AJ, Goldman AD. 2020. The complex phylogenetic relationships of a 4mC/6mA DNA methyltransferase in prokaryotes. Mol Phylogenet Evol.

Herman JJ, Sultan SE. 2016. DNA methylation mediates genetic variation for adaptive transgenerational plasticity. Proceedings of the Royal Society B: Biological Sciences 283.

Jeltsch A. 2003. Maintenance of species identity and controlling speciation of bacteria: A new function for restriction/modification systems? In: Gene.

Jiao J, Zhang B, Li ML, Zhang Z, Tian CF. 2022. The zinc-finger bearing xenogeneic silencer MucR in α-proteobacteria balances adaptation and regulatory integrity. ISME Journal.

Jones PA. 2012. Functions of DNA methylation: Islands, start sites, gene bodies and beyond. Nat Rev Genet.

Kahramanoglou C, Prieto AI, Khedkar S, Haase B, Gupta A, Benes V, Fraser GM, Luscombe NM, Seshasayee ASN. 2012. Genomics of DNA cytosine methylation in Escherichia coli reveals its role in stationary phase transcription. Nat Commun.

Kumar S, Karmakar BC, Nagarajan D, Mukhopadhyay AK, Morgan RD, Rao DN. 2018. N4-cytosine DNA methylation regulates transcription and pathogenesis in Helicobacter pylori. Nucleic Acids Res.

Ledermann R, Schulte CCM, Poole PS. 2021. How rhizobia adapt to the nodule environment. J Bacteriol.

Lercher MJ, Pál C. 2008. Integration of horizontally transferred genes into regulatory interaction networks takes many million years. Mol Biol Evol.

Letunic I, Bork P. 2021. Interactive tree of life (iTOL) v5: An online tool for phylogenetic tree display and annotation. Nucleic Acids Res.

Löytynoja A. 2014. Phylogeny-aware alignment with PRANK. Methods in Molecular Biology.

Militello KT, Mandarano AH, Varechtchouk O, Simon RD. 2014. Cytosine DNA methylation influences drug resistance in Escherichia coli through increased sugE expression. FEMS Microbiol Lett.

Mohapatra SS, Fioravanti A, Vandame P, Spriet C, Pini F, Bompard C, Blossey R, Valette O, Biondi EG. 2020. Methylation-dependent transcriptional regulation of crescentin gene (creS) by GcrA in Caulobacter crescentus. Mol Microbiol 114:127–139.

Oliveira PH. 2021. Bacterial Epigenomics: Coming of Age. mSystems.

Oliveira PH, Fang G. 2021. Conserved DNA Methyltransferases: A Window into Fundamental Mechanisms of Epigenetic Regulation in Bacteria. Trends Microbiol.

Oliveira PH, Touchon M, Rocha EPC. 2016. Regulation of genetic flux between bacteria by restriction-modification systems. Proc Natl Acad Sci U S A.

Page AJ, Cummins CA, Hunt M, et al. 2015. Roary: Rapid large-scale prokaryote pan genome analysis. Bioinformatics 31.22:3691–3693.

Pini F, De Nisco NJ, Ferri L, Penterman J, Fioravanti A, Brilli M, Mengoni A, Bazzicalupo M, Viollier PH, Walker GC, et al. 2015. Cell Cycle Control by the Master Regulator CtrA in Sinorhizobium meliloti. PLoS Genet.

Poole P, Ramachandran V, Terpolilli J. 2018. Rhizobia: From saprophytes to endosymbionts. Nat Rev Microbiol.

R Core Team. 2014. R Core Team (2014). R: A language and environment for statistical computing. R Foundation for Statistical Computing, Vienna, Austria. URL http://www.R-project.org/.

Reva ON, Swanevelder DZH, Mwita LA, Mwakilili AD, Muzondiwa D, Joubert M, Chan WY, Lutz S, Ahrens CH, Avdeeva L V., et al. 2019. Genetic, Epigenetic and Phenotypic Diversity of Four Bacillus velezensis Strains Used for Plant Protection or as Probiotics. Front Microbiol.

Riccardi C, Passeri I, Cangioli L, Fagorzi C, Mengoni A, Fondi M. 2022. MeStudio : crossing methylation and genomic features for comparative epigenomic analyses. :1–10.

Sánchez-Romero MA, Casadesús J. 2020. The bacterial epigenome. Nat Rev Microbiol.

Shaw LP, Rocha EPC, MacLean RC. 2022. Restriction-modification systems have shaped the evolution and distribution of plasmids across bacteria. bioRxiv [Internet]:2022.12.15.520556. Available from: https://www.biorxiv.org/content/10.1101/2022.12.15.520556v1

Shi WT, Zhang B, Li ML, Liu KH, Jiao J, Tian CF. 2022. The convergent xenogeneic silencer MucR predisposes α-proteobacteria to integrate AT-rich symbiosis genes. Nucleic Acids Res 50:8580–8598.

Stamatakis A. 2006. RAxML-VI-HPC: Maximum likelihood-based phylogenetic analyses with thousands of taxa and mixed models. Bioinformatics.

Touchon M, Hoede C, Tenaillon O, Barbe V, Baeriswyl S, Bidet P, Bingen E, Bonacorsi S, Bouchier C, Bouvet O, et al. 2009. Organised genome dynamics in the Escherichia coli species results in highly diverse adaptive paths. PLoS Genet.

Vasu K, Nagaraja V. 2013. Diverse Functions of Restriction-Modification Systems in Addition to Cellular Defense. Microbiology and Molecular Biology Reviews.

Yuan W, Zhang Y, Riaz L, Yang Q, Du B, Wang R. 2021. Multiple antibiotic resistance and DNA methylation in Enterobacteriaceae isolates from different environments. J Hazard Mater.

Zweiger G, Marczynski G, Shapiro L. 1994. A Caulobacter DNA methyltransferase that functions only in the predivisional cell. J Mol Biol.

