## Supplementary material for "The pan-epigenome of the symbiotic nitrogen fixing bacterium *Sinorhizobium meliloti* unravels unexpected variability of DNA-methylation profiles in closely related strains": Supplemenary information file

### Supplementary information

**Supplementary Dataset S1**. Genome coverage data and assignment of contigs to *S. meliloti* 1021 genome replicons. MS Excel file format (.xlsx file)

**Supplementary Dataset S2**. Folder containing summary pangenome statistics, core and accessory orthologs, alignments and R script. Zip archive

**Supplementary Dataset S3**. Number motifs and methylated motifs identified and their ratio. Data on number of motifs (RAW), number of methylated motifs (METH), fraction of methylated motifs (RATIO) related to functional features (CDSid, tIG and US sequences) and to replicons (CHR, chromosome; PA, pSymA; PB, pSymB and other) are reported. Red color in columns from "other" indicate missing data (i.e., absence of replicons other than chromosome, pSymA, pSymB). .xlsx file

#### *Putative replication initiation protein RepC coding genes*

**Table S1.** Locus tag (GenBank codes) of genes coding for RepC protein identified by genome annotation performed by the NCBI Prokaryotic Genome Annotation Pipeline (PGAP).

| **Strain/GenBank locus tags (repC gene)** | | | | | |
|---|---|---|---|---|---|
| 1A42 | I7G55_31090 | I7G55_31830 | | | |
| 4B41 | I7I49_19270 | I7I49_24670 | | | |
| 7B22 | I7G59_00015 | I7G59_20250 | I7G59_32180 | | |
| 8A52 | I7F98_01675 | I7F98_26050 | | | |
| AE608H | I7G86_26910 | I7G86_27090 | I7G86_35295 | | |
| AK58 | LOF18_00350 | LOF18_00570 | LOF18_12670 | LOF18_14870 | |
| AK75 | LOF12_24775 | LOF12_26095 | LOF12_32700 | | |
| AK83 | LOF17_00050 | LOF17_00235 | LOF17_14590 | LOF17_14705 | LOF17_15940 |

| | | | | | | |
|---|---|---|---|---|---|---|
| AL703GG | I7G01_00005 | I7G01_07565 | I7G01_13345 | I7G01_15370 | I7G01_15915 | I7G01_23550 |
| AL703H | I7G85_20290 | I7G85_24345 | I7G85_25215 | I7G85_25655 | | |
| AO641M | I7F95_03770 | I7F95_10950 | I7F95_11365 | I7F95_1954 | | |
| AO643DD | LOF23_19910 | LOF23_27580 | LOF23_35445 | | | |
| BL225C | LOF16_05515 | LOF16_15565 | | | | |
| CE480L | I7G00_19045 | I7G00_20425 | I7G00_34790 | | | |
| CL374FF | I7F97_00175 | I7F97_15100 | I7F97_15665 | I7F97_31915 | | |
| CO431A | LOF22_25585 | LOF22_33340 | | | | |
| CO438LL | LOF28_10065 | LOF28_10280 | LOF28_17945 | LOF28_18695 | | |
| H1 | I7G60_16970 | I7G60_17440 | I7G60_26575 | I7G60_27510 | | |
| NRG-185 | I7F13_00035 | I7F13_07680 | I7F13_15140 | I7F13_16300 | | |
| SM11 | LOF24_19230 | LOF24_26785 | LOF24_28185 | LOF24_28350 | | |
| T073 | I7F30_18610 | I7F30_19020 | I7F30_32420 | | | |

### *Methylated motifs identified*

The analysis of SMRT sequencing genome data with SMRT Link software ver. 8.0.0.80529 using default options identified 27 methylated motifs (Table S3A). Some of the motifs (shared colour) were further collapsed together since they share a common core motif (Table S3B).

**Table S2**. Raw (A) and collapsed (B) methylated motifs. Same colours indicated raw motifs collapsed into a single core motif.

A)

| Raw motifs |
|---|
| ACGGAG |
| BNNCGATCGV |
| BYCGATCG |
| CCCGGG |
| CGATCGV |
| CTCGAG |
| CTYCCAG |
| DCTGCAGGS |
| GANTC |
| GCCAGG |

| |
|---|
| GCCGGCH |
| GCCGGCYD |
| GCRDB |
| GNCGATCGVC |
| RAGCWGCTY |
| RCCAGCC |
| RCGATCGGC |
| RCTGCAGGS |
| RGATCY |
| SCTCGAG |
| TCGWCGA |
| TGGGCA |
| VGCCGGCCC |
| VNCGATCGV |
| YCGATCGD |
| YCGGCCGRV |
| YCTGCAG |

B)

| Motif | Type |
|---|---|
| ACGGAG | m6A |
| CGATCG | m6A |
| CCCGGG | m4C |
| CTCGAG | m6A |
| CTYCCAG | m6A |
| CTGCAG | m6A |
| GANTC | m6A |
| GCCAGG | m6A |
| GCCGGC | m4C |
| GCRDB | m4C |
| RAGCWGCTY | m4C |
| RCCAGCC | m4C |
| RGATCY | m6A |
| TCGWCGA | m4C |
| TGGGCA | m6A |
| YCGGCCGRV | m4C |

### *Putative DNA methyltransferases*

**Table S3.** Locus tag (GenBank codes) of DNA methyltransferases identified by genome annotation performed by the NCBI Prokaryotic Genome Annotation Pipeline (PGAP).

| **Strain/GenBank locus tags** | | | | | | | | | |
|---|---|---|---|---|---|---|---|---|---|
| 1A42 | I7G55_00850 | I7G55_04930 | I7G55_10975 | I7G55_10985 | I7G55_30135 | | | | |
| 4B41 | I7I49_01000 | I7I49_06525 | I7I49_06535 | I7I49_12810 | I7I49_16825 | I7I49_25660 | I7I49_29760 | | |
| 7B22 | I7G59_01700 | I7G59_05600 | I7G59_06395 | I7G59_06405 | I7G59_09770 | I7G59_19945 | I7G59_25705 | | |
| 8A52 | I7F98_02635 | I7F98_19595 | I7F98_23450 | I7F98_25655 | | | | | |
| AE608H | I7G86_05530 | I7G86_05560 | I7G86_07470 | I7G86_07480 | I7G86_13825 | I7G86_18415 | I7G86_29425 | I7G86_33635 | |
| AK58 | LOF18_01550 | LOF18_05945 | LOF18_13010 | LOF18_13015 | LOF18_16435 | LOF18_20780 | LOF18_26225 | LOF18_33095 | |
| AK75 | LOF12_00870 | LOF12_02240 | LOF12_04640 | LOF12_22340 | LOF12_26060 | | | | |
| AK83 | LOF17_06465 | LOF17_07730 | LOF17_21370 | LOF17_21380 | LOF17_29840 | LOF17_32675 | LOF17_34045 | | |
| AL703GG | I7G01_05265 | I7G01_08645 | I7G01_15395 | I7G01_27770 | I7G01_29440 | I7G01_32085 | | | |
| AL703H | I7G85_00865 | I7G85_04600 | I7G85_11135 | I7G85_32280 | | | | | |
| AO641M | I7F95_12345 | I7F95_16635 | I7F95_19520 | I7F95_20565 | I7F95_22230 | I7F95_24885 | I7F95_31755 | | |
| AO643DD | LOF23_01005 | LOF23_01035 | LOF23_07360 | LOF23_08780 | LOF23_11500 | LOF23_12275 | LOF23_18640 | LOF23_29880 | LOF23_34140 |
| BL225C | LOF16_06465 | LOF16_23090 | LOF16_24725 | LOF16_24735 | LOF16_28720 | LOF16_31075 | LOF16_32450 | | |
| CE480L | I7G00_06770 | I7G00_13630 | I7G00_17750 | I7G00_19200 | I7G00_21390 | | | | |
| CL374FF | I7F97_08515 | I7F97_12800 | I7F97_15640 | I7F97_16780 | I7F97_23625 | I7F97_26275 | I7F97_27930 | | |
| CO431A | LOF22_00865 | LOF22_02240 | LOF22_04595 | LOF22_08580 | LOF22_08590 | LOF22_10220 | LOF22_26565 | | |
| CO438LL | LOF28_11245 | LOF28_15495 | LOF28_20145 | LOF28_26550 | LOF28_29285 | LOF28_30775 | | | |
| H1 | I7G60_07150 | I7G60_12280 | I7G60_16015 | I7G60_24220 | | | | | |
| NRG185 | I7F13_06620 | I7F13_20825 | I7F13_21840 | I7F13_23250 | I7F13_23285 | I7F13_28155 | I7F13_31975 | | |
| SM11 | LOF24_12700 | LOF24_12730 | LOF24_14415 | LOF24_16980 | LOF24_18355 | LOF24_21550 | LOF24_25795 | LOF24_28160 | |
| T073 | I7F30_07070 | I7F30_07080 | I7F30_07745 | I7F30_11480 | I7F30_15215 | I7F30_31470 | | | |

### *Plasmid transfer experiment*

The artificial plasmid pMP7604 was used as probe for gene transfer experiments. Plasmid pMP7604 is a broad host range plasmid containing constitutively expressed mCherry gene and tetracycline resistance (Lagendijk et al., 2010). pMP7604 sequence is based on pME6031 vector (AddGene code V004776#) where the monomeric red fluorescent protein gene (GenBank accession number AY678264.1) downstream to a *tac* promoter (5'-TTGACAATTAATCATCGGCTCGTATAATG-3') have been cloned. The plasmid was inserted into *S. meliloti* strains by biparental mating using the rifampicin-resistant derivatives of *S. meliloti* strains as recipient strains, and *E. coli* S17-1 harboring pMP7604 as donor. The plasmid pMP7604 was then extracted from TY grown cultures of each *S. meliloti* strain with Qiagen miniprep kit. Quantity and quality of the plasmid DNA was estimated by 1% agarose gel electrophoresis in TAE buffer (40 mM Tris, 20 mM acetic acid, 1 mM EDTA). Then plasmid was extracted from each derivative *S. meliloti* strain culture and used in a pairwise combination to electroporate *S. meliloti* cells from the same and from the other 4 strains. DNA acquisition ratios were then computed as ratio between the efficiency of transformation (CFU/µg DNA) with not-self and with self-DNA. Preparation of electrocompetent *S. meliloti* cells vector was carried out as described in (Ferri et al., 2010). Briefly cells were grown in 100 ml TY medium at 30°C with shaking (130 rpm) until late exponential phase was reached. Cultures were cooled on ice for 15 minutes and centrifuged at 8000 rpm at 4°C for 10 minutes. Pellets were washed three times in ice cold MilliQ water and, finally, resuspended into 500 µl 10% glycerol to obtain 50 µl portions containing $1x10^8$ cells each. Electroporation was performed with $1x10^8$ *S. meliloti* competent cells and 1 µl vector (10-80 ng), according to the following settings: Voltage = 2.10 kV, Capacitance = 25 mF, Resistance = 200 Ohm.

In Table S3 the motifs identified in the *S. meliloti* genomes analyzed and present in pMP7604 are reported. The efficiency of transformation (CFU/µg DNA) with not-self and with self-DNA is reported in Table S4. Figure S1 reports the normalized efficiency of transformation against self-DNA (log10 scale) and Table S5 the p-values of pairwise contrasts.

**Figure S1**. **Pairwise transformation normalized efficiency.** Matrix of log10 fold change of DNA acquisition ratios against self-DNA (rows donors, columns recipient strains). Donors are indicated with the prefix "pMP7604_Me".

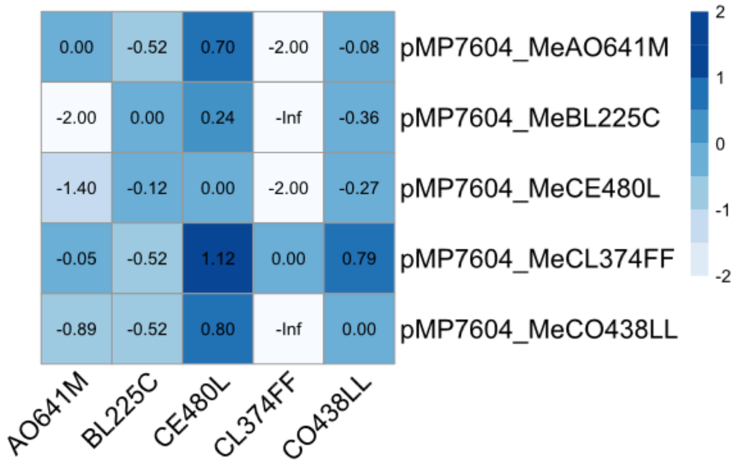

**Table S4**. Occurrences of motifs in the pMP7604 sequence.

| Motif | Occurrence in pMP7604 |
|---|---|
| ACGGAG | 2 |
| CGATCG | 3 |
| CCCGGG | 2 |
| CTCGAG | 1 |
| CTYCCAG | 2 |
| CTGCAG | 2 |
| GANTC | 24 |
| GCCAGG | 7 |
| GCCGGC | 28 |
| GCRDB | 258 |
| RAGCWGCTY | 0 |
| RCCAGCC | 3 |
| RGATCY | 7 |
| TCGWCGA | 1 |
| TGGGCA | 2 |
| YCGGCCGRV | 4 |

**Table S5**. **Gene transfer frequencies**. The table reports the details of the electroporation experiments performed. Number of recovered transformants and normalized efficiency of transformation (against self-DNA) are reported. Data are average of three replicate experiments. “p” indicates the donor (plasmid). The first two letters of strain codes are indicated.

| Donor | Recipient | Mean colonies | Mean efficiency |
|---|---|---|---|
| pAO | AO | 15.0 | 7.50E-09 |
| pBL | AO | 0.7 | 1.11E-10 |
| pCE | AO | 2.7 | 3.33E-10 |
| pCL | AO | 6.7 | 6.67E-09 |
| pCO | AO | 1.0 | 1.00E-09 |
| pAO | BL | 0.7 | 3.33E-10 |
| pBL | BL | 6.7 | 1.11E-09 |
| pCE | BL | 2.0 | 2.50E-10 |
| pCL | BL | 0.3 | 3.33E-10 |
| pCO | BL | 0.3 | 3.33E-10 |
| pAO | CE | 10.0 | 5.00E-09 |
| pBL | CE | 10.3 | 1.72E-09 |
| pCE | CE | 8.0 | 1.00E-09 |
| pCL | CE | 13.3 | 1.33E-08 |
| pCO | CE | 6.3 | 6.33E-09 |
| pAO | CL | 1.0 | 5.00E-10 |
| pBL | CL | 1.0 | 1.67E-10 |
| pCE | CL | 2.0 | 2.50E-10 |
| pCL | CL | 40.7 | 4.07E-08 |
| pCO | CL | 0.0 | 0.00E+00 |
| pAO | CO | 3.3 | 1.67E-09 |
| pBL | CO | 5.3 | 8.89E-10 |
| pCE | CO | 8.7 | 1.08E-09 |
| pCL | CO | 12.3 | 1.23E-08 |
| pCO | CO | 2.0 | 2.00E-09 |

**Table S6**. **Statistical significance of pairwise comparisons of transformation efficiencies**. Symmetric diagonal matrix of p-values from Dunn's post hoc test. Highly significant contrasts ($p<0.01$) are in bold. "p" indicates the donor (plasmid). The first two letters of strain codes are indicated.

| | pAO-AO | pBL-AO | pCE-AO | pCL-AO | pCO-AO | pAO-BL | pBL-BL | pCE-BL | pCL-BL | pCO-BL | pAO-CE | pBL-CE | pCE-CE | pCL-CE | pCO-CE | pAO-CL | pBL-CL | pCE-CL | pCL-CL | pCO-CL | pAO-CO | pBL-CO | pCE-CO | pCL-CO | pCO-CO |
|---|---|---|---|---|---|---|---|---|---|---|---|---|---|---|---|---|---|---|---|---|---|---|---|---|---|
| pAO-AO | | **0.004** | 0.019 | 0.918 | 0.077 | **0.009** | 0.185 | 0.013 | **0.009** | **0.009** | 0.778 | 0.452 | 0.150 | 0.763 | 0.910 | 0.036 | **0.007** | 0.013 | 0.541 | **0.002** | 0.332 | 0.112 | 0.218 | 0.778 | 0.259 |
| pBL-AO | **0.004** | | 0.579 | **0.005** | 0.259 | 0.771 | 0.116 | 0.686 | 0.771 | 0.771 | **0.009** | 0.032 | 0.145 | **0.001** | **0.005** | 0.424 | 0.836 | 0.686 | **0.000** | 0.785 | 0.054 | 0.191 | 0.096 | **0.001** | 0.077 |
| pCE-AO | 0.019 | 0.579 | | 0.025 | 0.566 | 0.792 | 0.309 | 0.880 | 0.792 | 0.792 | 0.039 | 0.112 | 0.366 | **0.008** | 0.026 | 0.807 | 0.728 | 0.880 | **0.003** | 0.408 | 0.169 | 0.452 | 0.267 | **0.009** | 0.225 |
| pCL-AO | 0.918 | **0.005** | 0.025 | | 0.096 | 0.012 | 0.221 | 0.017 | 0.012 | 0.012 | 0.858 | 0.516 | 0.181 | 0.686 | 0.993 | 0.046 | **0.010** | 0.017 | 0.475 | **0.002** | 0.387 | 0.137 | 0.259 | 0.700 | 0.305 |
| pCO-AO | 0.077 | 0.259 | 0.566 | 0.096 | | 0.402 | 0.658 | 0.469 | 0.402 | 0.402 | 0.137 | 0.309 | 0.742 | 0.038 | 0.098 | 0.742 | 0.356 | 0.469 | 0.017 | 0.161 | 0.424 | 0.858 | 0.592 | 0.040 | 0.522 |
| pAO-BL | **0.009** | 0.771 | 0.792 | 0.012 | 0.402 | | 0.201 | 0.910 | 1.000 | 1.000 | 0.020 | 0.064 | 0.243 | **0.004** | 0.013 | 0.611 | 0.933 | 0.910 | **0.001** | 0.572 | 0.102 | 0.309 | 0.169 | **0.004** | 0.140 |
| pBL-BL | 0.185 | 0.116 | 0.309 | 0.221 | 0.658 | 0.201 | | 0.243 | 0.201 | 0.201 | 0.296 | 0.566 | 0.910 | 0.104 | 0.225 | 0.440 | 0.172 | 0.243 | 0.053 | 0.065 | 0.721 | 0.792 | 0.925 | 0.108 | 0.843 |
| pCE-BL | 0.013 | 0.686 | 0.880 | 0.017 | 0.469 | 0.910 | 0.243 | | 0.910 | 0.910 | 0.027 | 0.082 | 0.292 | **0.005** | 0.017 | 0.693 | 0.843 | 1.000 | **0.002** | 0.498 | 0.127 | 0.366 | 0.207 | **0.005** | 0.172 |
| pCL-BL | **0.009** | 0.771 | 0.792 | 0.012 | 0.402 | 1.000 | 0.201 | 0.910 | | 1.000 | 0.020 | 0.064 | 0.243 | **0.004** | 0.013 | 0.611 | 0.933 | 0.910 | **0.001** | 0.572 | 0.102 | 0.309 | 0.169 | **0.004** | 0.140 |
| pCO-BL | **0.009** | 0.771 | 0.792 | 0.012 | 0.402 | 1.000 | 0.201 | 0.910 | 1.000 | | 0.020 | 0.064 | 0.243 | **0.004** | 0.013 | 0.611 | 0.933 | 0.910 | **0.001** | 0.572 | 0.102 | 0.309 | 0.169 | **0.004** | 0.140 |
| pAO-CE | 0.778 | **0.009** | 0.039 | 0.858 | 0.137 | 0.020 | 0.296 | 0.027 | 0.020 | 0.020 | | 0.638 | 0.247 | 0.560 | 0.866 | 0.069 | 0.016 | 0.027 | 0.371 | **0.004** | 0.492 | 0.191 | 0.342 | 0.572 | 0.397 |
| pBL-CE | 0.452 | 0.032 | 0.112 | 0.516 | 0.309 | 0.064 | 0.566 | 0.082 | 0.064 | 0.064 | 0.638 | | 0.492 | 0.292 | 0.522 | 0.178 | 0.053 | 0.082 | 0.172 | 0.016 | 0.829 | 0.402 | 0.631 | 0.301 | 0.707 |
| pCE-CE | 0.150 | 0.145 | 0.366 | 0.181 | 0.742 | 0.243 | 0.910 | 0.292 | 0.243 | 0.243 | 0.247 | 0.492 | | 0.082 | 0.185 | 0.510 | 0.211 | 0.292 | 0.040 | 0.083 | 0.638 | 0.880 | 0.836 | 0.085 | 0.756 |
| pCL-CE | 0.763 | **0.001** | **0.008** | 0.686 | 0.038 | **0.004** | 0.104 | **0.005** | **0.004** | **0.004** | 0.560 | 0.292 | 0.082 | | 0.679 | 0.016 | **0.003** | **0.005** | 0.756 | **0.001** | 0.204 | 0.059 | 0.125 | 0.985 | 0.153 |
| pCO-CE | 0.910 | **0.005** | 0.026 | 0.993 | 0.098 | 0.013 | 0.225 | 0.017 | 0.013 | 0.013 | 0.866 | 0.522 | 0.185 | 0.679 | | 0.047 | **0.010** | 0.017 | 0.469 | **0.002** | 0.392 | 0.140 | 0.263 | 0.693 | 0.309 |
| pAO-CL | 0.036 | 0.424 | 0.807 | 0.046 | 0.742 | 0.611 | 0.440 | 0.693 | 0.611 | 0.611 | 0.069 | 0.178 | 0.510 | 0.016 | 0.047 | | 0.553 | 0.693 | **0.007** | 0.283 | 0.259 | 0.611 | 0.387 | 0.017 | 0.332 |
| pBL-CL | **0.007** | 0.836 | 0.728 | **0.010** | 0.356 | 0.933 | 0.172 | 0.843 | 0.933 | 0.933 | 0.016 | 0.053 | 0.211 | **0.003** | **0.010** | 0.553 | | 0.843 | **0.001** | 0.631 | 0.085 | 0.271 | 0.145 | **0.003** | 0.118 |
| pCE-CL | 0.013 | 0.686 | 0.880 | 0.017 | 0.469 | 0.910 | 0.243 | 1.000 | 0.910 | 0.910 | 0.027 | 0.082 | 0.292 | **0.005** | 0.017 | 0.693 | 0.843 | | **0.002** | 0.498 | 0.127 | 0.366 | 0.207 | **0.005** | 0.172 |
| pCL-CL | 0.541 | **0.000** | **0.003** | 0.475 | 0.017 | **0.001** | 0.053 | **0.002** | **0.001** | **0.001** | 0.371 | 0.172 | 0.040 | 0.756 | 0.469 | **0.007** | **0.001** | **0.002** | | **0.000** | 0.114 | 0.028 | 0.065 | 0.742 | 0.082 |
| pCO-CL | **0.002** | 0.785 | 0.408 | **0.002** | 0.161 | 0.572 | 0.065 | 0.498 | 0.572 | 0.572 | **0.004** | 0.016 | 0.083 | **0.001** | **0.002** | 0.283 | 0.631 | 0.498 | **0.000** | | 0.028 | 0.114 | 0.053 | **0.001** | 0.041 |
| pAO-CO | 0.332 | 0.054 | 0.169 | 0.387 | 0.424 | 0.102 | 0.721 | 0.127 | 0.102 | 0.102 | 0.492 | 0.829 | 0.638 | 0.204 | 0.392 | 0.259 | 0.085 | 0.127 | 0.114 | 0.028 | | 0.535 | 0.792 | 0.211 | 0.873 |
| pBL-CO | 0.112 | 0.191 | 0.452 | 0.137 | 0.858 | 0.309 | 0.792 | 0.366 | 0.309 | 0.309 | 0.191 | 0.402 | 0.880 | 0.059 | 0.140 | 0.611 | 0.271 | 0.366 | 0.028 | 0.114 | 0.535 | | 0.721 | 0.061 | 0.645 |
| pCE-CO | 0.218 | 0.096 | 0.267 | 0.259 | 0.592 | 0.169 | 0.925 | 0.207 | 0.169 | 0.169 | 0.342 | 0.631 | 0.836 | 0.125 | 0.263 | 0.387 | 0.145 | 0.207 | 0.065 | 0.053 | 0.792 | 0.721 | | 0.130 | 0.918 |
| pCL-CO | 0.778 | **0.001** | **0.009** | 0.700 | 0.040 | **0.004** | 0.108 | **0.005** | **0.004** | **0.004** | 0.572 | 0.301 | 0.085 | 0.985 | 0.693 | 0.017 | **0.003** | **0.005** | 0.742 | **0.001** | 0.211 | 0.061 | 0.130 | | 0.158 |
| pCO-CO | 0.259 | 0.077 | 0.225 | 0.305 | 0.522 | 0.140 | 0.843 | 0.172 | 0.140 | 0.140 | 0.397 | 0.707 | 0.756 | 0.153 | 0.309 | 0.332 | 0.118 | 0.172 | 0.082 | 0.041 | 0.873 | 0.645 | 0.918 | 0.158 | |

### *Differences in methylation densities across coding and noncoding regions*

MeStudio procedure starts with matching the nucleotide motifs to the genomic sequence and map them to the corresponding category, which are extracted from the annotation file (GFF file). Categories are defined as: i) protein-coding genes with accordant (sense) strand (CDS), ii) regions that fall between annotated genes (true intergenic, tIG), iii) regions upstream to the reading frame of a gene, with accordant strand (US) (Figure S2). The CDS feature is defined by the ORF. The tIG is defined as the region between two different ORFs on both strands, as reported in Figure S2. The US region is defined, by default, as the portion of the genome between the end of an ORF and the beginning of the next one (Riccardi et al., 2023).

Results of statistical analysis of the overall densities of methylated motifs for CDS, tIG and US sequences in the *S. meliloti* replicons is reported in Table S7.

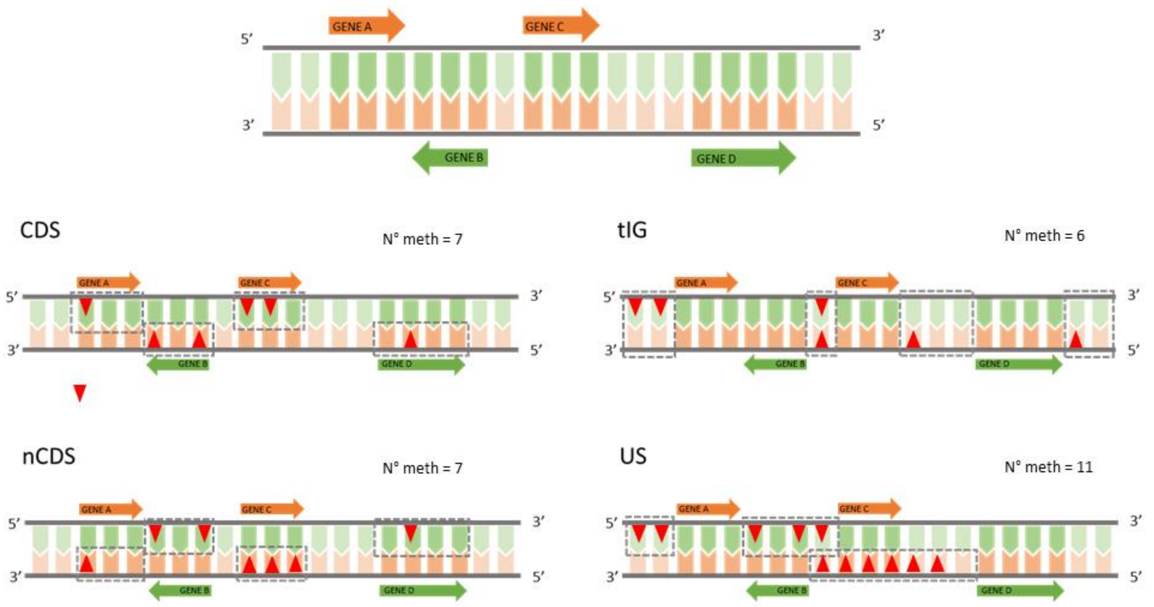

**Figure S2**. Graphical representation of the used terminology as in MeStudio software. CDS, coding sequence; tIG, intergenic sequence between two genes in opposite directions; US, upstream sequence to a coding sequence, (intergenic sequence between two genes having the same orientation).

**Table S7.** List of S. meliloti strains. The original isolation host (Medicago spp.) and country, genome length and number of contigs obtained are reported. All strains but H1 have been isolated from nodules (H1 from leaves)

| Strain code | Host plant (species, cultivar) | Isolation country | Genome lenght | Number of Contigs | Reference |
|---|---|---|---|---|---|
| 1A42 | *M. sativa "Hamadani"* | Iran | 6.973.268 | 4 | (Talebi et al., 2008) |
| 4B41 | *M. sativa "Nikshahri"* | Iran | 6.755.292 | 3 | (Talebi et al., 2008) |
| 7B22 | *M. sativa "Nikshahri"* | Iran | 6.791.724 | 4 | (Talebi et al., 2008) |
| 8A52 | *M. sativa "Hamadani"* | Iran | 6.735.207 | 3 | (Talebi et al., 2008) |
| AE608H | *M. sativa "Estival"* | Italy | 7.318.861 | 4 | (Carelli et al., 2000) |
| AK58 | *M. falcata* | Kazakhstan | 7.114.023 | 6 | (Roumiantseva et al., 2014) |
| AK75 | *M. lupulina* | Kazakhstan | 6.903.928 | 5 | (Roumiantseva et al., 2014) |

| | | | | | |
|---|---|---|---|---|---|
| AK83 | *M. falcata* | Kazakhstan | 7.171.952 | 7 | (Roumiantseva et al., 2014) |
| AL703GG | *M. sativa "Lodi"* | Italy | 7.487.501 | 9 | (Carelli et al., 2000) |
| AL703H | *M. sativa "Lodi"* | Italy | 7.020.338 | 5 | (Carelli et al., 2000) |
| AO641M | *M. sativa "Oneida"* | Italy | 7.503.659 | 7 | (Carelli et al., 2000) |
| AO643DD | *M. sativa "Oneida"* | Italy | 7.442.138 | 4 | (Carelli et al., 2000) |
| BL225C | *M. sativa "Lodi"* | Italy | 6.996.973 | 3 | (Carelli et al., 2000) |
| CE480L | *M. sativa "Estival"* | Italy | 7.340.765 | 5 | (Carelli et al., 2000) |
| CL374FF | *M. sativa "Lodi"* | Italy | 7.436.312 | 6 | (Carelli et al., 2000) |
| CO431A | *M. sativa "Oneida"* | Italy | 7.001.362 | 3 | (Carelli et al., 2000) |
| CO438LL | *M. sativa "Oneida"* | Italy | 6.709.320 | 6 | (Carelli et al., 2000) |
| H1 | *M. sativa* | Italy | 6.912.818 | 6 | (Galardini et al., 2013) |
| NGR185 | *M. sativa* | Canada | 7.329.821 | 7 | (Prevost et al., 1999) |
| SM11 | *M. sativa* | Germany | 7.502.918 | 5 | (Stiens et al., 2006) |
| T073 | *M. truncatula* | Tunisia | 6.958.496 | 4 | (Porter & Simms, 2014) |

### References

Carelli, M., Gnocchi, S., Fancelli, S., Mengoni, A., Paffetti, D., Scotti, C., & Bazzicalupo, M. (2000). Genetic Diversity and Dynamics of Sinorhizobium meliloti Populations Nodulating Different Alfalfa Cultivars in Italian Soils. In *APPLIED AND ENVIRONMENTAL MICROBIOLOGY* (Vol. 66, Issue 11). https://journals.asm.org/journal/aem

Ferri, L., Gori, A., Biondi, E. G., Mengoni, A., & Bazzicalupo, M. (2010). Plasmid electroporation of Sinorhizobium strains: The role of the restriction gene hsdR in type strain Rm1021. *Plasmid*. https://doi.org/10.1016/j.plasmid.2010.01.001

Galardini, M., Pini, F., Bazzicalupo, M., Biondi, E. G., & Mengoni, A. (2013). Replicon-dependent bacterial genome evolution: The case of Sinorhizobium meliloti. *Genome Biology and Evolution*. https://doi.org/10.1093/gbe/evt027

Lagendijk, E. L., Validov, S., Lamers, G. E. M., De Weert, S., & Bloemberg, G. V. (2010). Genetic tools for tagging Gram-negative bacteria with mCherry for visualization in vitro and in natural habitats, biofilm and pathogenicity studies. *FEMS Microbiology Letters*. https://doi.org/10.1111/j.1574-6968.2010.01916.x

Porter, S. S., & Simms, E. L. (2014). Selection for cheating across disparate environments in the legume-rhizobium mutualism. *Ecology Letters*, *17*(9), 1121–1129. https://doi.org/10.1111/ele.12318

Prevost, D., Drouin, P., & Antoun, H. (1999). The potential use of cold-adapted rhizobia to improve symbiotic nitrogen fixation in legumes cultivated in temperate regions. In R. Margesin & F. Schinner (Eds.), *Biotechnological applications of cold-adapted organisms* (pp. 161–176).

Riccardi, C., Passeri, I., Cangioli, L., Fagorzi, C., Fondi, M., & Mengoni, A. (2023). Crossing Bacterial Genomic Features and Methylation Patterns with MeStudio: An Epigenomic Analysis Tool. *International Journal of Molecular Sciences*, *24*(1). https://doi.org/10.3390/ijms24010159

Riccardi, C., Passeri, I., Cangioli, L., Fagorzi, C., Mengoni, A., & Fondi, M. (2022). *MeStudio : crossing methylation and genomic features for comparative epigenomic analyses*. 1–10.

Roumiantseva, M. L., Muntyan, V. S., Mengoni, A., & Simarov, B. V. (2014). ITS-polymorphism of salt-tolerant and salt-sensitive native isolates of Sinorhizoblum meliloti-symbionts of alfalfa, clover and fenugreek plants. *Russian Journal of Genetics*, *50*(4), 348–359. https://doi.org/10.1134/S1022795414040103

Stiens, M., Schneiker, S., Keller, M., Kuhn, S., Pühler, A., & Schlüter, A. (2006). Sequence analysis of the 144-kilobase accessory plasmid pSmeSM11a, isolated from a dominant Sinorhizobium meliloti strain identified during a long-term field release experiment. *Applied and Environmental Microbiology*, *72*(5), 3662–3672. https://doi.org/10.1128/AEM.72.5.3662-3672.2006

Talebi, M. B., Bahar, M., Saeidi, G., Mengoni, A., & Bazzicalupo, M. (2008). Diversity of Sinorhizobium strains nodulating Medicago sativa from different Iranian regions. *FEMS Microbiology Letters*, *288*(1), 40–46. https://doi.org/10.1111/j.1574-6968.2008.01329.x
